# Metabolic insight into bacterial community assembly across ecosystem boundaries

**DOI:** 10.1101/758615

**Authors:** Nathan I. Wisnoski, Mario E. Muscarella, Megan L. Larsen, Ariane L. Peralta, Jay T. Lennon

## Abstract

The movement of organisms across habitat boundaries has important consequences for populations, communities, and ecosystems. However, because most species are not well adapted to all habitat types, dispersal into suboptimal habitats could induce physiological changes associated with persistence strategies that influence community assembly. For example, high rates of cross-boundary dispersal are thought to maintain sink populations of terrestrial bacteria in aquatic habitats, but these bacteria may also persist by lowering their metabolic activity, introducing metabolic heterogeneity that buffers the population against niche selection. To differentiate between these assembly processes, we analyzed bacterial composition along a hydrological flow path from terrestrial soils through an aquatic reservoir by sequencing the active and total (active + inactive) portions of the community. When metabolic heterogeneity was ignored, our data were consistent with views that cross-boundary dispersal is important for structuring aquatic bacterial communities. In contrast, we found evidence for strong niche selection when metabolic heterogeneity was explicitly considered, suggesting that, relative to persistence strategies, dispersal may have a weaker effect on aquatic community assembly than previously thought. By accounting for metabolic heterogeneity in complex communities, our findings clarify the roles of local- and regional-scale assembly processes in terrestrial-aquatic meta-ecosystems.

## INTRODUCTION

The movement of material and energy across habitat boundaries is important for the structure and function of recipient ecosystems (Polis et al. 2004, Gounand et al. 2018a). These spatial resource subsidies can stabilize population dynamics, alter food web structure, and modify biogeochemical cycles (Polis et al. 2004, Massol et al. 2011). However, in complex landscapes linked by spatial fluxes of resources and organisms, the process of community assembly remains less clear (Gounand et al. 2018a). Meta-ecosystem theory predicts that poorly adapted species dispersed across ecosystem boundaries will be eliminated from the recipient habitat via niche selection (Massol et al. 2017, Gounand et al. 2018a), unless resource flows sufficiently homogenize the landscape (Gravel et al. 2010). However, if generalist species are capable of tolerating a range of environmental conditions, then cross-boundary dispersal could affect community assembly in recipient habitats (Haegeman and Loreau 2014).

Habitats at the terrestrial-freshwater interface are ideal for addressing questions about meta-ecosystem ecology (Gounand et al. 2018b). Terrestrial ecosystems export large quantities of organic matter that support aquatic food webs, often through bacterial pathways (Berggren et al. 2010). Furthermore, many of the bacteria responsible for processing allochthonous subsidies in aquatic habitats may be derived from terrestrial ecosystems via coupled transport with resource flows (Ruiz-González et al. 2015b). For example, in some north temperate lakes, it is estimated that nearly 10^20^ bacterial cells are transported annually from terrestrial to aquatic ecosystems (Bergström and Jansson 2000). These high immigration rates should influence the composition and activity of bacterial assemblages via metacommunity processes, such as source-sink dynamics or mass effects that overcome niche selection (Crump et al. 2012, Lindström and Langenheder 2012, Ruiz-González et al. 2015a).

Although cross-boundary flows have been well documented, the fate of terrestrial-derived bacteria in aquatic ecosystems remains unclear (Langenheder and Lindström 2019). In part, this may be because both dispersal- and selection-based perspectives overlook the range of metabolic states within microbial communities. In nature, some microorganisms may respond to favorable environmental conditions via rapid growth, while others may face challenging conditions that limit or prevent growth (Lever et al. 2015). Many bacteria have evolved persistence strategies (e.g., spores, cysts, resting stages, slow growth) that buffer against harsh environmental transitions, such as those encountered when dispersed along terrestrial-aquatic flow paths (Barcina et al. 1997, Lennon and Jones 2011). By weakening the strength of local niche selection relative to dispersal (Nemergut et al. 2013, Locey et al. 2019, Wisnoski et al. 2019), these persistence strategies may increase the apparent similarity between terrestrial and aquatic bacterial communities, especially when techniques are used that lend equal weight to active, slow growing, and dormant bacteria (e.g., 16S rRNA gene sequencing). As a result, the importance of terrestrial-derived bacteria in aquatic community assembly may be obscured when inferred from diversity patterns that do not explicitly consider the metabolic heterogeneity that exists within bacterial communities.

In this study, we explored microbial community assembly along a hydrological flow path of a small reservoir. In this type of system, inputs from the terrestrial landscape occur upstream in the riverine zone, directional surface flow orients the passive dispersal of bacteria through the lacustrine zone, and emigration occurs over an impoundment (Thornton et al. 1990; Fig. 1). We hypothesized that dispersal maintains terrestrial-derived bacteria in the reservoir, promoting local (α) diversity and homogenizing among-site (β) diversity at the aquatic-terrestrial interface, but that these taxa may not be metabolically active. Owing to niche selection, we hypothesized that only a subset of the immigrating terrestrial bacteria become metabolically active members of the aquatic community.

**Fig. 1.**
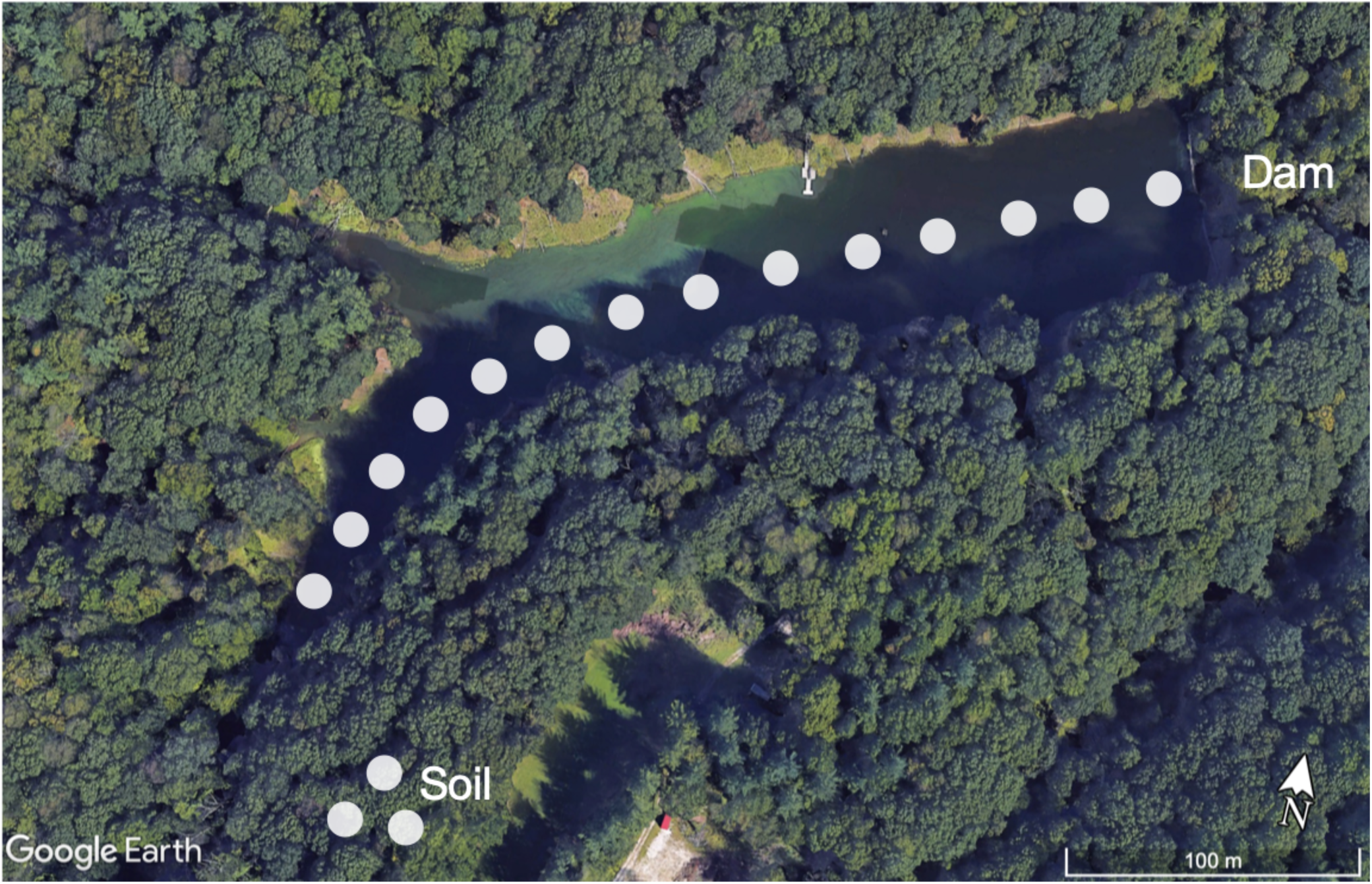
University Lake, Indiana University Research and Teaching Preserve, Bloomington, Indiana, USA. Points indicate sampling locations along the terrestrial-aquatic transect, from upstream soils, through the stream inlet, across the lacustrine zone, and over the dam. Image source: Google Earth.

## METHODS

### Study system

University Lake is a meso-eutrophic reservoir located in Griffy Woods, Bloomington, Indiana, USA (39.189, −86.503) (Fig. 1). Created in 1911, the 3.2 ha impoundment has an operating volume of 150,000 m^3^. With a maximum depth of 10 m, University Lake is fed by three streams that drain mature oak-beech-maple forest. The underlying geology is Harrodsburg limestone on ridgetops and Borden siltstone/shale in valleys. The thin unglaciated soils surrounding the reservoir are Brownstown-Gilwood silt loams.

### Bacterial community structure

We collected surface-water samples along a longitudinal transect through University Lake in July 2013, filtering epilimnetic biomass from 200 mL of water onto 0.2 µm Supor Filters (47 mm diameter, Pall). We characterized composition of the active and total portions of the bacterial communities by sequencing 16S rRNA genes (DNA) and transcripts (RNA), respectively. While sequences recovered from the DNA pool can come from active or inactive individuals, sequences from the RNA pool are commonly used to study active microorganisms given that rRNA transcripts have short half-lives and that ribosomes are required by growing cells for protein synthesis (Molin and Givskov 1999, Steiner et al. 2019, Bowsher et al. 2019, Locey et al. 2019). Sequences were processed in *mothur* (v. 1.41.1, Schloss et al. 2009) and operational taxonomic units (OTUs) were created using the OptiClust algorithm (Westcott and Schloss 2017). See supplement for detailed methods.

### Quantifying patterns of diversity along the flow path

We analyzed within sample (α) and among sample (β) diversity along the flow path. We estimated α-diversity using rarefaction in the ‘iNEXT’ R package (Hsieh et al. 2016), following singleton-correction for sequence data (Chiu and Chao 2016). Hill numbers for a given order, *q*, were used to weigh common and rare species using the equation 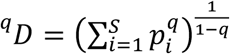, where *p*_*i*_ is the relative abundance of species *i* = 1, …, *S*. The value ^*q*^*D* is the number of equally abundant species that would yield the observed value of a diversity metric, such as richness (*q* = 0), Shannon’s index (*q* = 1), or Simpson’s index (*q* = 2). We measured β-diversity as average percent similarity (1 – Bray-Curtis) between each sample using the ‘vegan’ package in R (Oksanen et al. 2019). We used indicator-variables multiple regression to test for the main effects and interaction of molecule type (RNA vs. DNA) and flow-path distance on α- and β-diversity.

To make inferences about niche selection on terrestrial bacteria, we measured changes in the relative abundances of OTUs that were recovered in the DNA and RNA pools. To quantify the loss rate of terrestrial-derived bacteria, we used the slope of a simple linear regression between distance along the transect and the number of terrestrial OTUs present but never active in aquatic samples. To determine possible contributions of soil-derived taxa to active aquatic diversity, we focused on OTUs that were metabolically active across a majority of aquatic samples (we present results for a 75% threshold in main text, others in supplement). All statistical analyses were conducted in R (version 3.5.2, R Core Team 2018).

## RESULTS

Patterns of bacterial diversity along the flow path were strongly influenced by metabolic heterogeneity (Fig. 2a, *R*^*2*^ *=* 0.84, p < 0.001), as shown by significant differences in slope and intercept captured by the indicator variable (Table 1). In the total aquatic bacterial community (DNA), richness was highest near the terrestrial-aquatic interface and declined toward the dam. In comparison, the active aquatic richness was lower and less variable along the transect. Differences in α-diversity between active and total portions of the community were highest near the terrestrial-aquatic interface (e.g., there were 78% fewer taxa in the active subset, Table 1). Diversity differences were greatest when rare and common taxa were equally weighted (*q* = 0), as might be expected if immigrant or dormant taxa are rare. When dominant taxa were weighted more heavily (*q* = 1, 2), the active portion remained less diverse, but the decay rates of diversity became indistinguishable between the two portions of the community (Table 1).

**Table 1.** Linear model coefficients of active and total α-diversity along the transect examined at different levels of *q*, which represents equal weighting of rare and common taxa (q = 0), proportional weighting (q = 1), and biased weighting toward common taxa (q = 2). In these models, intercepts represent estimates of diversity at each order near the terrestrial-aquatic interface, with the RNA term capturing the reduced diversity in the active subset. With increasing order, the distance × RNA interaction becomes weaker, signifying that diversity decays at similar rates in the active and total communities as common taxa are increasingly weighted.

| Order ($q$) | Diversity | Term | Estimate | Std. Error | Statistic | p-value |
| --- | --- | --- | --- | --- | --- | --- |
| 0 | Richness | Intercept | 1497 | 100.6 | 14.88 | $<10^{-4}$ |
| 0 | Richness | Distance | -3.176 | .4976 | -6.381 | $<10^{-4}$ |
| 0 | Richness | RNA | -1170 | 142.3 | -8.222 | $<10^{-4}$ |
| 0 | Richness | Distance × RNA | 2.985 | .7003 | 4.263 | .0003 |
| 1 | Shannon | Intercept | 153.7 | 19.41 | 7.921 | $<10^{-4}$ |
| 1 | Shannon | Distance | -.2941 | 0.096 | -3.062 | 0.0053 |
| 1 | Shannon | RNA | -123.9 | 27.46 | -4.513 | .0001 |
| 1 | Shannon | Distance × RNA | 0.2457 | .1352 | 1.818 | 0.0815 |
| 2 | Simpson | Intercept | 55.44 | 6.47 | 8.57 | $<10^{-4}$ |
| 2 | Simpson | Distance | -.0783 | .032 | -2.446 | 0.0221 |
| 2 | Simpson | RNA | -36.78 | 9.151 | -4.019 | 0.0005 |
| 2 | Simpson | Distance × RNA | .0402 | .045 | 0.8918 | 0.3813 |

**Fig. 2.**
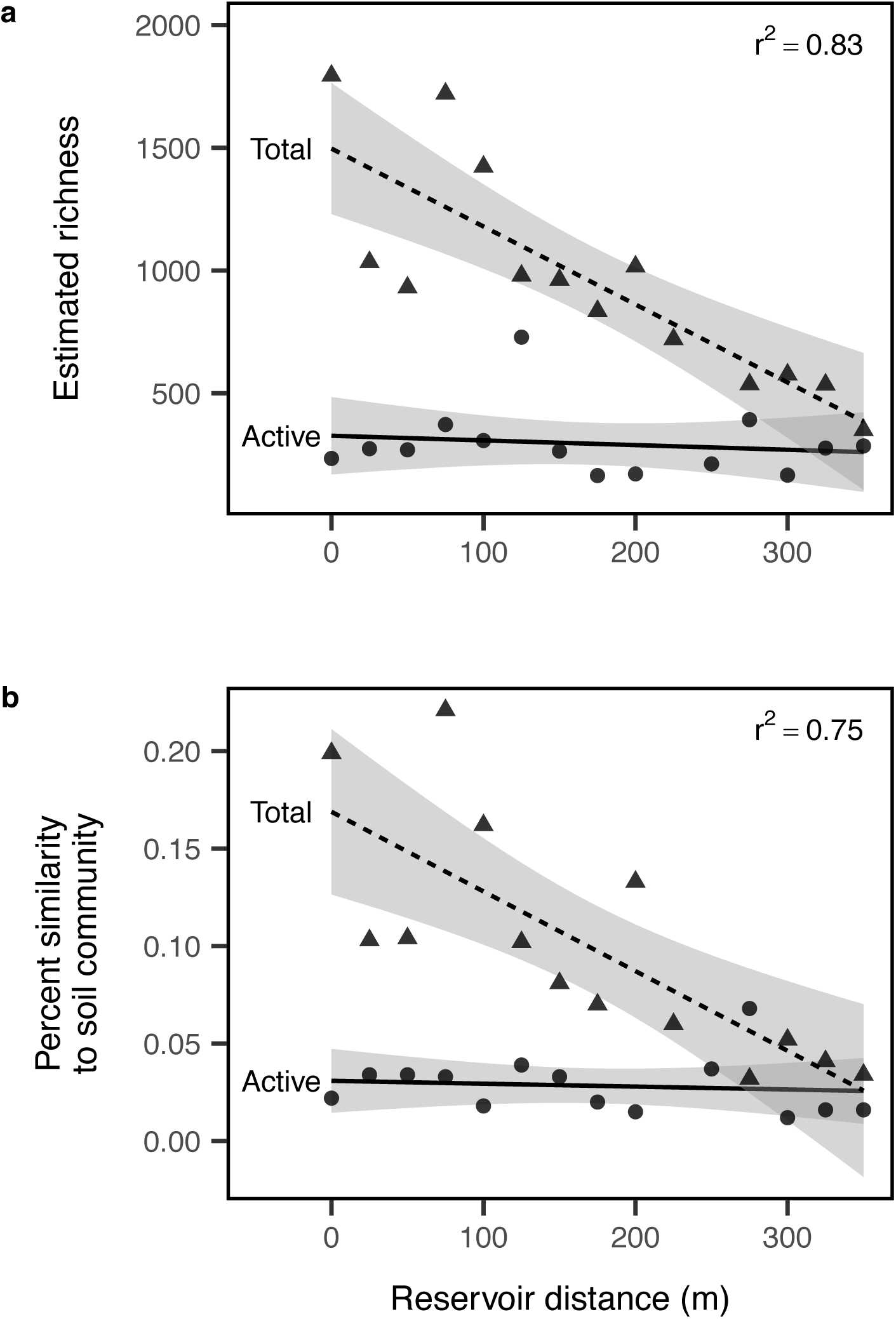
Terrestrial influence on aquatic microbial diversity. (a) Estimated alpha diversity (richness, ^*1*^*D*) in the active (light gray circles) and total (dark gray triangles) aquatic communities along the reservoir transect. (b) The average percent similarity to the soil samples for active and total aquatic communities declines with distance away from the terrestrial-aquatic interface (0 m).

Metabolic heterogeneity also has strong effects on β-diversity (Fig. 2b). Similarity between terrestrial soil and aquatic samples was highest near the terrestrial-aquatic interface and decreased toward the dam (*R*^*2*^ = 0.75, p < 0.001). However, maximum similarity to soils and the rate of decay in similarity differed between the total and active portion of the community. Near the stream inlet, similarity to soils was nearly 6 times higher in the total community than in the active portion (*Intercept* = 0.169, β_RNA_ = −0.138), and similarity to soils declined linearly toward the dam (β_distance_ = −4 × 10^−4^, β_distance × RNA_ = −3.9×10^−4^). In contrast, the active portion remained dissimilar to terrestrial soils along the entire transect (Fig. 2b).

We detected a small number of habitat generalists, but most terrestrial taxa did not appear to colonize the aquatic community. The majority of taxa present in both soil and aquatic communities were never detected in any active aquatic sample (∼88% of taxa remained inactive), which accounted for roughly 4.5% of all reads in the total reservoir community. The richness of these taxa declined exponentially (first-order decay, *k* = 2.57 × 10^−3^=, *r*^*2*^=0.81, p < 0.001) with distance from the stream inlet (Fig. 3a). However, 8% of taxa present in soils were detected at least once in the active aquatic community. Of the soil-derived taxa detected in at least 75% of active aquatic samples, 18 declined along the transect, but 11 were maintained at high relative abundances in the active aquatic community (Fig. 3; see supplement for list of taxa).

**Fig. 3.**
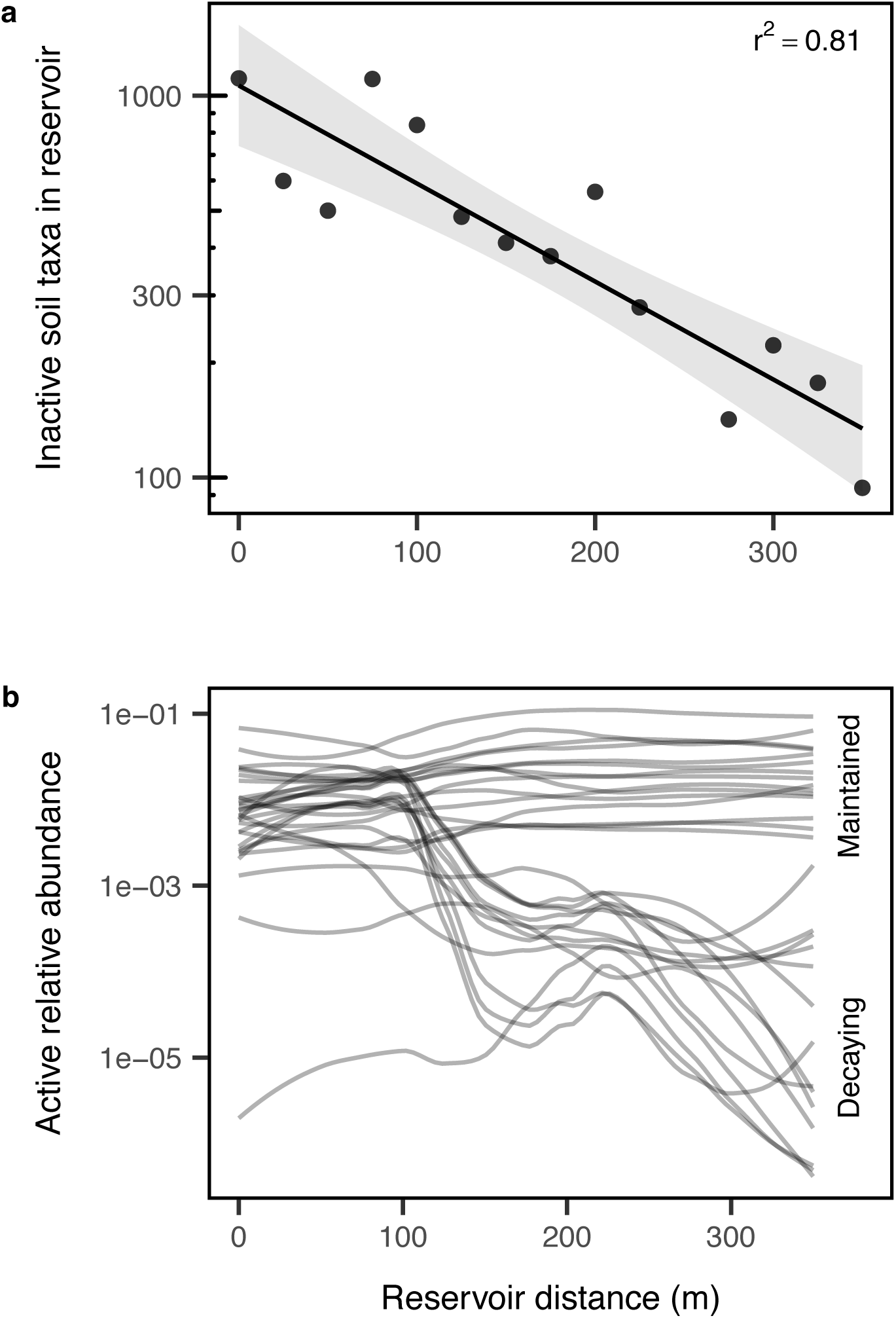
Fate of terrestrial-derived taxa in the reservoir. (a) Number of taxa detected in soils but never detected in active aquatic samples declines exponentially away from the terrestrial-aquatic interface with a first-order decay constant *k* = 2.57 × 10^−3^=. (b) Taxa detected in at least 75% of active aquatic samples either decay in abundance along the transect or are maintained. We used local polynomial regression (LOESS) to visualize relative abundances for each OTU along the transect.

## DISCUSSION

Our results demonstrate that the importance of dispersal for community assembly across ecosystem boundaries depends on the metabolic activity of dispersers in the meta-ecosystem. Along a terrestrial-aquatic flow path, the influence of terrestrial bacteria on aquatic bacterial α- and β-diversity was highest near the terrestrial-aquatic interface. This pattern, consistent with terrestrial immigration playing an important role in aquatic community assembly (i.e., mass effects), was only detected when metabolic heterogeneity was not considered. In contrast, α- diversity and similarity to soils were substantially lower in the metabolically active portion of the aquatic community (Table 1; Fig. 2), suggesting a hidden role for niche selection in the aquatic habitat that was only apparent when incorporating metabolic information. In fact, most terrestrial-derived taxa were not detected in the active aquatic community and decayed exponentially away from the terrestrial-aquatic interface (Fig. 3a). Altogether, our findings are consistent with our hypotheses that most terrestrial-derived taxa fail to colonize aquatic habitats and that only a small number of habitat generalists may be able to colonize aquatic environments from nearby terrestrial landscapes. Our study also suggests the potentially overlooked role of metabolic heterogeneity in spatially heterogeneous metacommunities and meta-ecosystems.

### Metabolic heterogeneity informs aquatic community assembly

Inferring community assembly processes from diversity patterns is challenging because species can be present in a habitat for reasons other than habitat suitability (e.g., high dispersal, persistence traits). Accounting for metabolic heterogeneity helps distinguish favorable from suboptimal habitats by detecting the responses of actively growing organisms (e.g., Muscarella et al. 2016), providing insight into the fate and potential functions of dispersers in recipient ecosystems. The frequent detection of terrestrial bacteria in aquatic ecosystems has elevated the role of dispersal for structuring aquatic diversity, but our results suggest that local aquatic environments can still impose harsh biotic or abiotic filters on the metabolically active subset of the aquatic community (Fig. 2). Thus, the strength of niche selection against terrestrial-derived bacteria in aquatic habitats may increase with metabolic activity levels of cross-boundary dispersers.

### Exponential decay of soil-derived bacteria in aquatic ecosystems

Dispersing across an ecosystem boundary is likely a harsh transition for many bacteria. Although most active aquatic taxa were also detected in nearby soils, only a minority of taxa present in soils were common in the active aquatic community (Fig. 3). Consistent with previous terrestrial-aquatic meta-ecosystem studies (Mariadassou et al. 2015, Monard et al. 2016), our results suggest that active abundance is highest in preferred habitat types. The exponential decay of metabolically inactive terrestrial taxa away from the terrestrial-aquatic interface also resembles diversity declines near river margins (Power et al. 2004). This exponential loss could be due to physical factors (e.g., settling or volumetric dilution) or biotic interactions (e.g., consumption, competition, or lysis following reactivation) that are not offset by reproduction. Future studies that differentiate activities at a finer resolution (e.g., slow growing, dormant with the potential to reactivate, or even dead) (Carini et al. 2016, Lennon et al. 2018) could further illuminate the fate of cross-boundary dispersers in meta-ecosystems. In general, the exponential decay suggests that terrestrial influences on aquatic bacterial diversity may be localized near ecosystem boundaries.

Nevertheless, a subset of taxa detected in soils were active in the aquatic community. Some became less common along the transect, which could reflect a riverine-to-lacustrine environmental gradient, or a reduction in mass effects (Fig. 3b). These decaying taxa included representatives from the Actinobacteria (*Arthrobacter, Micrococcus, Solirubrobacter*), Bacteroidetes (*Flavobacterium, Pedobacter*), Proteobacteria (α: *Bradyrhizobium, Sphingomonas*; β: *Duganella, Comamonas*; and γ: *Pseudomonas* sp.), some of which are abundant and ubiquitous in soils (Delgado-Baquerizo et al. 2018). In contrast, taxa maintained in the active aquatic community may have wide niche breadths allowing them to be habitat generalists, or they may be of aquatic origin (e.g., dispersed by floods, animals, or wind, but our soil sampling locations were chosen to minimize this possibility). These potential habitat generalists included taxa belonging to the Actinomycetales, Bacteroidetes (order Sphingobacteriales), Proteobacteria (α: order Rhizobiales, β: family Comamonadaceae, γ: *Acinetobacter*), and Verrucomicrobia (class Spartobacteria). In sum, most terrestrial-derived bacteria may possess persistence strategies that allow them to persist on the periphery of aquatic ecosystems, but habitat generalists that cross ecosystem boundaries could influence aquatic bacterial community assembly.

### Metabolic heterogeneity in metacommunities and meta-ecosystems

Our work provides empirical evidence that accounting for metabolic heterogeneity may improve our understanding of metacommunity and meta-ecosystem processes (Massol et al. 2017, Wisnoski et al. 2019). Cross-boundary dispersal can expose organisms to harsh environmental conditions, against which they may be buffered through metabolic flexibility (e.g., slow growth, dormancy). While generalists may be able to colonize a range of habitat types in meta-ecosystems (Haegeman and Loreau 2014), specialist dispersal may require coupling with resource subsidies or persistence strategies that buffer against suboptimal conditions. Metabolically explicit community assembly also has implications for ecosystem functioning in a spatial context. While high dispersal is predicted to impede ecosystem functioning by creating species-environment mismatches (Leibold et al. 2017), these effects may be reduced if dispersers are metabolically inactive. Thus, metabolic heterogeneity may be an important link between individuals, communities, and ecosystems across spatial scales.

## ACKNOWLEDGEMENTS

We thank Nick Nelson and the Holland Summer Science Program, the National Science Foundation DEB-1442246 (JTL), and the US Army Research Office Grant W911NF-14-1-0411 (JTL). Data and code can be found at NCBI (BioProject PRJNA547598) and GitHub (https://github.com/LennonLab/ReservoirGradient).

## REFERENCES

Barcina, I., P. Lebaron, and J. Vives-Rego. 1997. Survival of allochthonous bacteria in aquatic systems: a biological approach. FEMS Microbiology Ecology 23:1–9.

Berggren, M., L. Ström, H. Laudon, J. Karlsson, A. Jonsson, R. Giesler, A.-K. Bergström, and M. Jansson. 2010. Lake secondary production fueled by rapid transfer of low molecular weight organic carbon from terrestrial sources to aquatic consumers. Ecology Letters 13:870–880.

Bergström, A.-K., and M. Jansson. 2000. Bacterioplankton production in humic Lake Örträsket in relation to input of bacterial cells and input of allochthonous organic carbon. Microbial Ecology 39:101–115.

Bowsher, A. W., P. J. Kearns, and A. Shade. 2019. 16S rRNA/rRNA gene ratios and cell activity staining reveal consistent patterns of microbial activity in plant-associated soil. mSystems 4:e00003–19.

Carini, P., P. J. Marsden, J. W. Leff, E. E. Morgan, M. S. Strickland, and N. Fierer. 2016. Relic DNA is abundant in soil and obscures estimates of soil microbial diversity. Nature Microbiology 2:1–6.

Chiu, C. H., and A. Chao. 2016. Estimating and comparing microbial diversity in the presence of sequencing errors. PeerJ 4:e1634.

Crump, B. C., L. A. Amaral-Zettler, and G. W. Kling. 2012. Microbial diversity in arctic freshwaters is structured by inoculation of microbes from soils. The ISME journal 6:1629–1639.

Delgado-Baquerizo, M., A. M. Oliverio, T. E. Brewer, A. Benavent-González, D. J. Eldridge, R. D. Bardgett, F. T. Maestre, B. K. Singh, and N. Fierer. 2018. A global atlas of the dominant bacteria found in soil. Science 359:320–325.

Gounand, I., E. Harvey, C. J. Little, and F. Altermatt. 2018a. Meta-ecosystems 2.0: rooting the theory into the field. Trends in Ecology & Evolution 33:36–46.

Gounand, I., C. J. Little, E. Harvey, and F. Altermatt. 2018b. Cross-ecosystem carbon flows connecting ecosystems worldwide. Nature Communications 9:4825.

Gravel, D., F. Guichard, M. Loreau, and N. Mouquet. 2010. Source and sink dynamics in meta-ecosystems. Ecology 91:2172–2184.

Haegeman, B., and M. Loreau. 2014. General relationships between consumer dispersal, resource dispersal and metacommunity diversity. Ecology Letters 17:175–184.

Hsieh, T. C., K. H. Ma, and A. Chao. 2016. iNEXT: an R package for rarefaction and extrapolation of species diversity (Hill numbers). Methods in Ecology and Evolution 7:1451–1456.

Langenheder, S., and E. S. Lindström. 2019. Factors influencing aquatic and terrestrial bacterial community assembly. Environmental Microbiology Reports 11:306–315.

Leibold, M. A., J. M. Chase, and S. K. M. Ernest. 2017. Community assembly and the functioning of ecosystems: how metacommunity processes alter ecosystems attributes. Ecology 98:909–919.

Lennon, J. T., and S. E. Jones. 2011. Microbial seed banks: the ecological and evolutionary implications of dormancy. Nature Reviews Microbiology 9:119–130.

Lennon, J. T., M. E. Muscarella, S. A. Placella, and B. K. Lehmkuhl. 2018. How, when, and where relic DNA affects microbial diversity. mBio 9:e00637–18.

Lever, M. A., K. L. Rogers, K. G. Lloyd, J. Overmann, B. Schink, R. K. Thauer, T. M. Hoehler, and B. B. Jørgensen. 2015. Life under extreme energy limitation: a synthesis of laboratory- and field-based investigations. FEMS Microbiology Reviews 39:688–728.

Lindström, E. S., and S. Langenheder. 2012. Local and regional factors influencing bacterial community assembly. Environmental Microbiology Reports 4:1–9.

Locey, K. J., M. E. Muscarella, M. L. Larsen, S. R. Bray, S. E. Jones, and J. T. Lennon. 2019. Dormancy dampens the microbial distance-decay relationship. bioRxiv:717546.

Mariadassou, M., S. Pichon, and D. Ebert. 2015. Microbial ecosystems are dominated by specialist taxa. Ecology Letters 18:974–982.

Massol, F., F. Altermatt, I. Gounand, D. Gravel, M. A. Leibold, and N. Mouquet. 2017. How life-history traits affect ecosystem properties: effects of dispersal in meta-ecosystems. Oikos 126:532–546.

Massol, F., D. Gravel, N. Mouquet, M. W. Cadotte, T. Fukami, and M. A. Leibold. 2011. Linking community and ecosystem dynamics through spatial ecology. Ecology Letters 14:313–323.

Molin, S., and M. Givskov. 1999. Application of molecular tools for in situ monitoring of bacterial growth activity. Environmental Microbiology 1:383–391.

Monard, C., S. Gantner, S. Bertilsson, S. Hallin, and J. Stenlid. 2016. Habitat generalists and specialists in microbial communities across a terrestrial-freshwater gradient. Scientific Reports 6:37719.

Muscarella, M. E., S. E. Jones, and J. T. Lennon. 2016. Species sorting along a subsidy gradient alters bacterial community stability. Ecology 97:2034–2043.

Nemergut, D. R. et al. 2013. Patterns and processes of microbial community assembly. Microbiology and Molecular Biology Reviews 77:342–356.

Oksanen, J. et al. 2019. vegan: Community Ecology Package. R package version 2.5-4. https://CRAN.R-project.org/package=vegan.

Polis, G. A., M. E. Power, and G. R. Huxel, editors. 2004. Food webs at the landscape level. The University of Chicago Press, Chicago, IL.

Power, M. E. et al. 2004. River-to-watershed subsidies in an old-growth conifer forest. Pages 217–240 in G. A. Polis, M. E. Power, and G. R. Huxel, editors. Food webs at the landscape level. The University of Chicago Press, Chicago, IL.

R Core Team. 2018. R: A language and environment for statistical computing. R Foundation for Statistical Computing, Vienna, Austria. URL https://www.R-project.org/.

Ruiz-González, C., J. P. Niño-García, and P. A. del Giorgio. 2015a. Terrestrial origin of bacterial communities in complex boreal freshwater networks. Ecology Letters 18:1198–1206.

Ruiz-González, C., J. P. Niño-García, J.-F. Lapierre, and P. A. del Giorgio. 2015b. The quality of organic matter shapes the functional biogeography of bacterioplankton across boreal freshwater ecosystems. Global Ecology and Biogeography 24:1487–1498.

Schloss, P. D. et al. 2009. Introducing mothur: Open-source, platform-independent, community-supported software for describing and comparing microbial communities. Applied and Environmental Microbiology 75:7537–7541.

Steiner, P. A., D. D. Corte, J. Geijo, C. Mena, T. Yokokawa, T. Rattei, G. J. Herndl, and E. Sintes. 2019. Highly variable mRNA half-life time within marine bacterial taxa and functional genes. Environmental Microbiology.

Thornton, K. W., B. L. Kimmel, and F. E. Payne, editors. 1990. Reservoir limnology: ecological perspectives. John Wiley & Sons, New York, NY.

Westcott, S. L., and P. D. Schloss. 2017. OptiClust, an improved method for assigning amplicon-based sequence data to operational taxonomic units. mSphere 2:e00073–17.

Wisnoski, N. I., M. A. Leibold, and J. T. Lennon. 2019. Dormancy in metacommunities. The American Naturalist 194:135–151.

