## Supplementary File for "Metabolic insight into bacterial community assembly across ecosystem boundaries"

### SUPPLEMENTAL METHODS

Surface water samples were obtained approximately every 25 m along a longitudinal transect from the lacustrine zone near the dam to the two major streams feeding University Lake (Fig. 1). We used a Quanta Hydrolab (OTT, Kempton, Germany) water sonde to measure temperature, dissolved oxygen, pH, and conductivity of the epilimnion at each site. We collected water samples at each site for biological and chemical analyses. We measured total phosphorus (TP) concentrations using the ammonium molybdate method (Wetzel and Likens 2000).

*Sample preparation* — For aquatic samples, we extracted total nucleic acids (RNA and DNA) from the filters using the MoBio PowerWater RNA extraction kit and the DNA elution accessory kit (Carlsbad, CA) and cleaned the extracts via ethanol precipitation. We treated RNA extracts with DNase (Invitrogen) to degrade DNA prior to cDNA synthesis via the SuperScript III First Strand Synthesis Kit and random hexamer primers (Invitrogen). For soil samples, we extracted DNA with the PowerSoil DNA isolation kit (MoBio, Carlsbad, CA). Once DNA and cDNA samples were cleaned and quantified, we amplified the 16S rRNA gene (DNA) and transcript (cDNA) using barcoded primers (515F and 806R) targeting the V4 hypervariable region (Caporaso et al. 2012). We purified sequence libraries using the AMPure XP purification kit (Bechman), quantified using the Quant-it PicoGreen dsDNA kit (Invitrogen), and pooled at equal molar ratios (final concentration:10 ng per library). After pooling, we sequenced the libraries on the Illumina MiSeq platform using 250 × 250 bp paired end reads (Reagent Kit v2) at the Indiana University Center for Genomics and Bioinformatics Sequencing Facility. Paired-end raw 16S rRNA sequences reads were assembled into contigs, quality-trimmed, and aligned to the Silva Database (version 132) (Quast et al. 2013). Chimeric sequences were detected and removed using the VSEARCH algorithm (Rognes et al. 2016). We created OTUs by first splitting the sequences based on the RDP taxonomy (Cole et al. 2009), and then binning sequences in to operational taxonomic units (OTUs) based on 97% sequence similarity. All initial sequence processing was completed using the software package *mothur* (version 1.41.1, Schloss et al. 2009).

### TABLES

Table S1 – Taxa found in soils that get rarer along the transect (declining).

| OTU | slope | pval | Phylum | Class | Order | Family | Genus |
|---|---|---|---|---|---|---|---|
| Otu00009 | -5.159e-05 | 0.02755 | Proteobacteria | Gammaproteobacteria | Pseudomonadales | Pseudomonadaceae | Pseudomonas |
| Otu00010 | -4.34e-05 | 0.5521 | Proteobacteria | Proteobacteria_unclassified | Proteobacteria_unclassified | Proteobacteria_unclassified | Proteobacteria_unclassified |
| Otu00011 | -1.949e-05 | 0.6012 | Proteobacteria | Betaproteobacteria | Betaproteobacteria_unclassified | Betaproteobacteria_unclassified | Betaproteobacteria_unclassified |
| Otu00018 | -4.676e-05 | 0.02114 | Proteobacteria | Gammaproteobacteria | Pseudomonadales | Pseudomonadaceae | Pseudomonas |
| Otu00022 | -2.524e-05 | 0.1182 | Verrucomicrobia | Opitutae | Opitutae_unclassified | Opitutae_unclassified | Opitutae_unclassified |
| Otu00028 | -3.068e-05 | 0.02359 | Proteobacteria | Gammaproteobacteria | Pseudomonadales | Pseudomonadaceae | Pseudomonas |
| Otu00030 | -2.244e-06 | 0.2763 | Actinobacteria | Actinobacteria | Actinomycetales | Micrococcaceae | Micrococcus |
| Otu00039 | -8.596e-06 | 0.1787 | Proteobacteria | Betaproteobacteria | Burkholderiales | Comamonadaceae | Comamonas |
| Otu00045 | -8.037e-06 | 0.5276 | Proteobacteria | Betaproteobacteria | Burkholderiales | Oxalobacteraceae | Oxalobacteraceae_unclassified |
| Otu00059 | -6.541e-05 | 0.02553 | Actinobacteria | Actinobacteria | Actinomycetales | Micrococcaceae | Arthrobacter |
| Otu00065 | -5.579e-05 | 0.02116 | Bacteroidetes | Sphingobacteriia | Sphingobacteriales | Sphingobacteriaceae | Pedobacter |
| Otu00072 | -1.895e-05 | 0.09149 | Proteobacteria | Alphaproteobacteria | Sphingomonadales | Sphingomonadaceae | Sphingomonas |
| Otu00077 | -5.886e-05 | 0.01187 | Bacteroidetes | Flavobacteriia | Flavobacteriales | Flavobacteriaceae | Flavobacterium |
| Otu00086 | -1.265e-05 | 0.03621 | Proteobacteria | Alphaproteobacteria | Rhizobiales | Bradyrhizobiaceae | Bradyrhizobium |
| Otu00094 | -2.23e-05 | 0.03169 | Proteobacteria | Betaproteobacteria | Burkholderiales | Oxalobacteraceae | Duganella |
| Otu00095 | -3.578e-05 | 0.03614 | Proteobacteria | Betaproteobacteria | Burkholderiales | Comamonadaceae | Comamonadaceae_unclassified |
| Otu00170 | -2.494e-05 | 0.02878 | Bacteroidetes | Sphingobacteriia | Sphingobacteriales | Sphingobacteriaceae | Sphingobacteriaceae_unclassified |
| Otu00545 | -1.236e-06 | 0.02985 | Actinobacteria | Actinobacteria | Solirubrobacterales | Solirubrobacteraceae | Solirubrobacter |

Table S2 – Taxa found in soils that get more common along the transect (maintained).

| OTU | slope | pval | Phylum | Class | Order | Family | Genus |
|---|---|---|---|---|---|---|---|
| Otu00001 | 1.436e-05 | 0.07999 | Proteobacteria | Betaproteobacteria | Burkholderiales | Comamonadaceae | Comamonadaceae_unclassified |
| Otu00002 | 0.0002115 | 0.002237 | Actinobacteria | Actinobacteria | Actinomycetales | Actinomycetales_unclassified | Actinomycetales_unclassified |
| Otu00003 | 9.899e-05 | 0.006441 | Verrucomicrobia | Spartobacteria | Spartobacteria_unclassified | Spartobacteria_unclassified | Spartobacteria_unclassified |
| Otu00005 | 3.61e-05 | 0.01737 | Bacteroidetes | Sphingobacteriia | Sphingobacteriales | Chitinophagaceae | Sediminibacterium |
| Otu00006 | 6.575e-06 | 0.1618 | Bacteroidetes | Sphingobacteriia | Sphingobacteriales | Saprospiraceae | Saprospiraceae_unclassified |
| Otu00012 | 7.541e-06 | 0.09905 | Proteobacteria | Betaproteobacteria | Burkholderiales | Comamonadaceae | Comamonadaceae_unclassified |
| Otu00014 | 8.464e-05 | 0.0007891 | Actinobacteria | Actinobacteria | Actinomycetales | Actinomycetales_unclassified | Actinomycetales_unclassified |
| Otu00023 | 3.267e-07 | 0.8 | Proteobacteria | Gammaproteobacteria | Pseudomonadales | Moraxellaceae | Acinetobacter |
| Otu00029 | 3.32e-05 | 0.004456 | Actinobacteria | Actinobacteria | Actinomycetales | Actinomycetales_unclassified | Actinomycetales_unclassified |
| Otu00032 | 3.56e-06 | 0.8341 | Bacteroidetes | Bacteroidetes_unclassified | Bacteroidetes_unclassified | Bacteroidetes_unclassified | Bacteroidetes_unclassified |
| Otu00033 | 9.129e-06 | 0.7085 | Proteobacteria | Alphaproteobacteria | Rhizobiales | Rhizobiales_unclassified | Rhizobiales_unclassified |

**FIGURES**

Figure S1 — Environmental variables across the transect fit with a loess smoother.

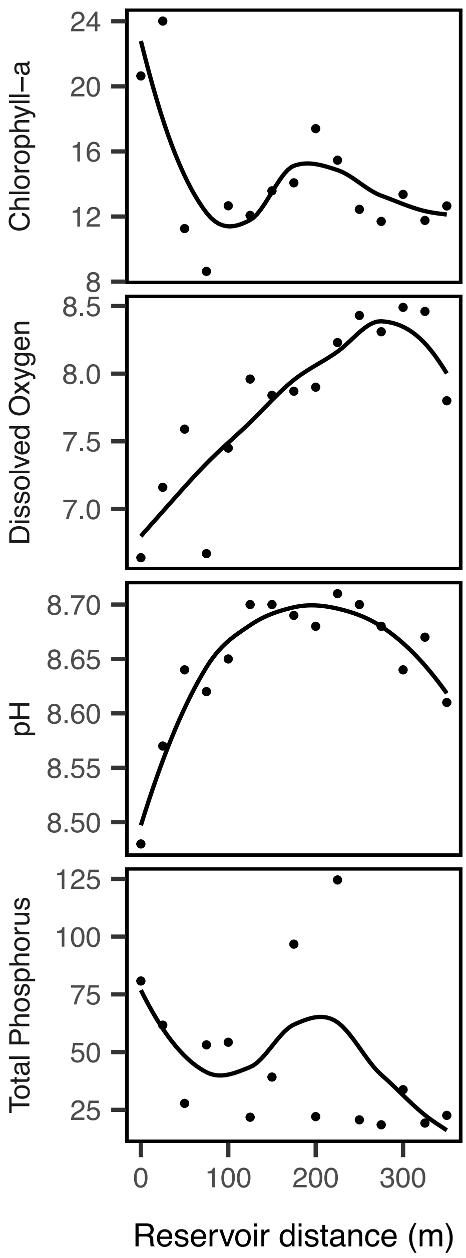

Figure S2 – Sensitivity of terrestrial-derived OTU fate to threshold of OTU incidence cutoff (minimum fraction of sites detected). We present cutoff of 0.75 in the main text, but qualitative conclusions remain consistent across thresholds, with some taxa declining and others maintained along the gradient.

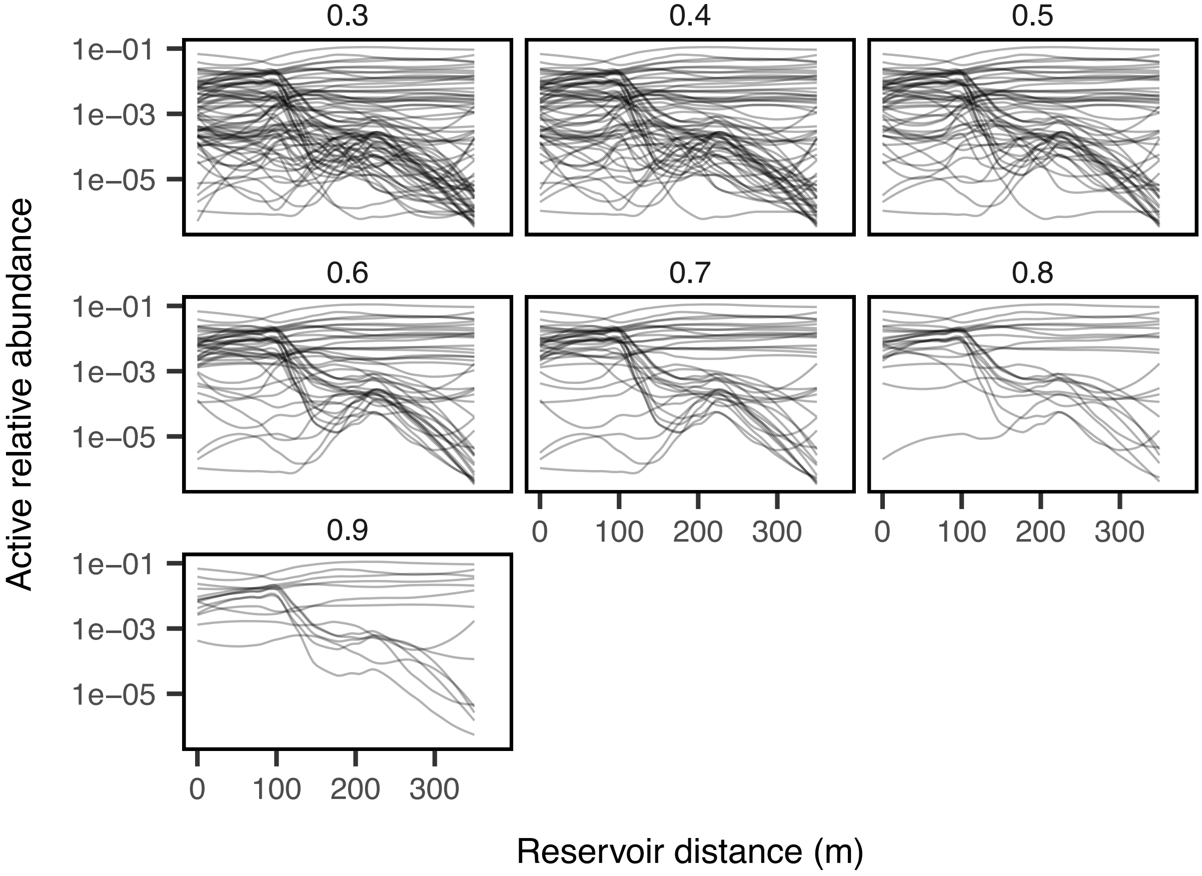

### REFERENCES


Caporaso, J. G., C. L. Lauber, W. A. Walters, D. Berg-Lyons, J. Huntley, N. Fierer, S. M. Owens, J. Betley, L. Fraser, M. Bauer, N. Gormley, J. A. Gilbert, G. Smith, and R. Knight. 2012. Ultra-high-throughput microbial community analysis on the Illumina HiSeq and MiSeq platforms. The ISME Journal 6:1621–1624.

Cole, J. R., Q. Wang, E. Cardenas, J. Fish, B. Chai, R. J. Farris, A. S. Kulam-Syed-Mohideen, D. M. McGarrell, T. Marsh, G. M. Garrity, and J. M. Tiedje. 2009. The Ribosomal Database Project: Improved alignments and new tools for rRNA analysis. Nucleic Acids Research 37:141–145.

Quast, C., E. Pruesse, P. Yilmaz, J. Gerken, T. Schweer, P. Yarza, J. Peplies, and F. O. Glöckner. 2013. The SILVA ribosomal RNA gene database project: improved data processing and web-based tools. Nucleic Acids Research 41:590–596.

Rognes, T., T. Flouri, B. Nichols, C. Quince, and F. Mahé. 2016. VSEARCH: a versatile open source tool for metagenomics. PeerJ 4:e2584–e2584.

Schloss, P. D., S. L. Westcott, T. Ryabin, J. R. Hall, M. Hartmann, E. B. Hollister, R. A. Lesniewski, B. B. Oakley, D. H. Parks, C. J. Robinson, J. W. Sahl, B. Stres, G. G. Thallinger, D. J. Van Horn, and C. F. Weber. 2009. Introducing mothur: Open-source, platform-independent, community-supported software for describing and comparing microbial communities. Applied and Environmental Microbiology 75:7537–7541.

Wetzel, R. G., and G. E. Likens. 2000. Limnological analyses. 3rd ed. Springer, New York, NY.
